# Two distinct integrin binding sites on MMP9 drive cancer invasion by mediating integrin membrane trafficking & stabilization

**DOI:** 10.1101/2024.04.01.587542

**Authors:** Sarbajeet Dutta, Soumili Sarkar, Simran Tolani, Asrafuddoza Hazari, Shamik Sen

**Affiliations:** Dept. of Biosciences & Bioengineering, IIT Bombay, Mumbai 400076

**Keywords:** MMP9, invasion, integrin β1 (ITGβ1), matrix degradation, non-proteolytic function

## Abstract

Matrix stiffening has been established to drive cancer progression through increased activity of matrix metalloproteases (MMPs) which degrade the matrix creating paths for migration. However, the non-proteolytic functions of MMPs in cancer invasion remain relatively less understood. Here we have probed the importance of proteolytic and non-proteolytic functions of MMP9, which exhibits robust stiffness dependent expression and secretion in highly invasive cancer cells. We show that while MMP9 sustains spreading and 2D migration non-proteolytically by stabilizing focal adhesions, MMP9 proteolytic activity is essential for 3D invasion. We then establish the function of two distinct integrin β1 (ITG β1) binding sites on MMP9, with the hemopexin domain mediating co-packaging and co-transport of ITG β1/MMP9 to the cell periphery, and the RGD domain stabilizing ITG β1 on the cell membrane prior to matrix degradation. Together, our results illustrate how MMP9 optimizes cancer invasion by spatiotemporally integrating matrix remodeling with adhesion formation.

## Introduction

The extracellular matrix (ECM) constitutes the acellular component of all tissues [1], [2], [3]. Cells residing within the tissue deposit a variety of protein/non-protein components to form this complex 3D structure. Although ECM homeostasis is strictly regulated in normal tissues, this homeostasis is compromised in cancerous tissues [4]. Excess deposition of fibrillar collagen and crosslinking by enzymes such as lysyl oxidase leads to matrix stiffening in multiple epithelial cancers including breast cancer and pancreatic cancer [5], [6]. Altered cell-matrix signaling is associated with malignant transformation leading to epithelial-to-mesenchymal transition (EMT) [7] through increased secretion of matrix degrading enzymes such as matrix metalloproteinases (MMPs). Indeed, expression of multiple MMPs including MMP1, 2, 3, 7, 9, 13, and 14 are closely associated with cancer invasion and worse prognosis [8], [9], [10], [11], [12].

The multidomain structural architecture of MMPs with different substrate specificities is well documented in the literature. While MMP1, 8, 13, and 18 mostly act on fibrillar collagen, MMP2 and 9 can also degrade non-fibrillar collagen and gelatin [13][14]. Despite the structural similarities in their catalytic domain, pro-peptide, and signal peptide, individual MMPs have distinct structural traits that define their functional properties. While most membrane bound MMPs have a transmembrane/cytoplasmic domain (MMP14, 15, 16, and 24), MMP17 and 25 have GPI anchoring to the membrane. Except for MMP7 and 26, all MMPs have hemopexin repeats; in addition, MMP2/9 possess fibronectin type II repeats in their catalytic domains, which can purportedly bind to gelatin, collagen types I and IV, and laminin [14].

The proteolytic activity of MMPs mediated by their catalytic domain has been studied extensively and is well established. For example, overexpression of MMP3 and 14 has been shown to induce breast cancer and melanoma [15], [16], whereas MMP2 and 14 promote cell invasion by cleaving laminin 5 [17], [18]. MMP3 and 7 degrade E-cadherin in MCF7 and MDCK cells, causing cancer cells to separate from the tissue [19]. A recent study has implicated MMP9 in tumor progression in triple-negative breast cancer [20]. MMP9 expression increases during the early stages of melanoma development [21]. In the animal model, inhibiting MMP9 lowers lung metastases [22].

Beyond their proteolytic function, a growing body of literature has identified the importance of non-proteolytic activity of MMPs mediated by their hemopexin, transmembrane, and cytoplasmic domains, via interactions with diverse cell surface proteins that aid in invasion and cancer growth. According to emerging evidence in the literature, MMP9 and 14 can localize on the cell membrane via a variety of mechanisms. MMP14 interacts with CD44, a hyaluronan receptor, and promotes invasion by releasing CD44 from the cell membrane in pancreatic and breast cancer [23]. MMP14 has also been shown to engage integrin β1 (ITGβ1) via its transmembrane/cytoplasmic domain and activate MAPK in mouse mammary epithelial cells leading to branching morphogenesis [24]. The interaction of CD44 with MMP9 via the hemopexin domain leads to enhanced invasion and angiogenesis in mouse mammary carcinoma and human melanoma cells [25]. In melanoma cells, MMP9 has also been shown to interact with ITGβ1 via its hemopexin domain, leading to the formation of cell-substrate adhesions [26], [27].

While insightful, these studies have not parsed the proteolytic and non-proteolytic functions of MMP9 in cancer invasion. We address this question by first showing that among several MMPs, MMP9 exhibits strong stiffness dependent expression and secretion in highly invasive cancer cells. Cell rounding and loss of cell motility upon inhibition of MMP proteolytic activity are attributed to defects in adhesion stability [28]. While similar effects were observed upon MMP9 knockdown, restoration of cell spreading and 2D motility in knockdown cells re-expressing catalytically inactive MMP9 through stabilization of adhesions, but no restoration of invasion in 3D collagen matrices, suggests that MMP9 non-proteolytically regulates adhesion formation, with matrix degradation essential for 3D invasion. We then show that two ITGβ1 binding sites, a RGD motif in the catalytic domain and a previously identified peptide sequence in the hemopexin domain [29], collectively regulate adhesion formation. While MMP9/ ITGβ1 association via the hemopexin domain is necessary for integrin trafficking to the membrane and MMP9 secretion, the RGD motif is responsible for stabilizing ITGβ1 on the cell membrane. Collectively, our study illustrates how invasiveness is optimally sustained by MMP9 through a combination of path generation via proteolytic remodeling and stabilization of integrins at the leading edge.

## Materials & Methods

### TCGA Data analysis

The data from The Cancer Genome Atlas (TCGA) was obtained through curation of the UCSC Xena Browser, an online resource accessible at https://xenabrowser.net/. The analysis focused exclusively on the TCGA breast cancer (BRCA) and TCGA lung cancer (LUNG) datasets, among the various types of cancer data available. The gene expression levels of MMP1, MMP2, MMP9, and MMP14 were determined by RNA sequencing (Illumina HiSeq) in both cancerous and normal tissue samples. Subsequently, the data obtained was organized and subjected to analysis. A total of 1097 individuals diagnosed with breast cancer and 114 healthy individuals were included in the breast cancer analysis. Similarly, the lung cancer analysis comprised 1017 individuals diagnosed with lung cancer and 110 healthy individuals.

### Cell culture & reagents

In our study, the following human cell lines were used: HT1080 invasive fibrosarcoma cell line, MDA-MB-231 invasive breast cancer cell line, A549 lung carcinoma cell line, MCF-7 breast cancer cell line, MCF-10A breast epithelial cell line, and Normal Human Dermal Fibroblasts (HDF) cells isolated from the healthy human tissue sample. The cancer cells utilized in this study were procured from the NCCS Cell Repository in Pune. The HEK293T cells utilized for lentivirus generation were generously provided by the laboratory of Dr. Prakriti Tayalia. Cells were cultured in Dulbecco’s Modified Eagle Medium (DMEM) (Cat # 12100046, Gibco) supplemented with 10% fetal bovine serum (FBS) (Cat. # 10270106, Gibco) and 1% antibiotic (Cat. # A002, HiMedia) under 5% CO_2_ conditions. For drug treatment experiments, GM6001 (Cat. # 2983, Tocris Bioscience) and SB-3CT (Cat. # 6088, Tocris Bioscience) were used at a working concentration of 10 µM [30] and ARP100 (Cat. # 2621, Tocris Bioscience) was used at a working concentration 50nM [31].

### Cloning and generation of stable cell lines

MMP9 full-length cDNA was cloned from HT1080 cDNA to create expression plasmids using gene-specific cloning primers M9_F_KpnI and M9_R_NotI (Supplementary Table 1). The amplified product was first cloned into pcDNA3 (Addgene plasmid# 13031), followed by MMP9-fused eGFP that was subcloned into the pLenti-puro vector (Addgene plasmid# 39481) to generate the MMP9 full-length expression plasmid (M9FL) using primer pairs M9_F_EcoRI and eGFP_R_XbaI. To generate a catalytic inactive mutant (Δcat), the primer pairs M9_E4A_F and M9_E4A_R were used to introduce a point mutation of glutamic acid (E) to alanine (A) at amino acid position 402 (E402A) in M9FL. RGE, ΔB1, and ΔB1Δcat plasmids were obtained by amplifying M9FL and M9Δcat plasmids with the primer pairs M9_RGE_F and R and M9_B1_F and R, respectively (Supplementary Table 1).

Stable MMP9 knockdown cell lines were generated using two independent shRNA sequences (CCACAACATCACCTATTGGAT and CAGTTTCCATTCATCTTCCAA; Sigma Aldrich). The transduced cells were selected using puromycin (Catalogue number CMS8861, HiMedia). Re-expression of M9FL and other MMP9 cDNAs was done using lentiviral transduction, produced by HEK293T cells according to the standard protocol [32]. Cells were transduced with lentiviral particles in the presence of polybrene (8 g/mL, Cat# TR-1003-G, Merck); after 48 hours of transduction, positive cells were sorted using fluorescent-assisted cell sorting (FACS).

### Cell Experiments on Polyacrylamide (PA) gels

PA gels were prepared as per previously established lab protocols to generate 0.5 kPa soft gels mimicking the stiffness of normal breast tissue and 5 kPa malignant tissue [30]. Gels were functionalized with 0.1M sulfo-succinimidyl 6-(4’-azido-2’-nitrophenylamino) hexanoate (Sulfo-SANPAH) (Cat# 22589, Thermo Scientific) dissolved in HEPES before overnight incubation in collagen type-I solution (Catalogue # A1048301, Thermo Scientific) at 4^0^C to achieve a theoretical coating density of 10 *μ*g/cm^2^. For experiments, cells were seeded sparsely (2000 cells/cm^2^) and cultured for 24 hours before imaging and motility experiments; cells were seeded at high density (20000 cells/cm^2^) for RT-PCR, western blotting, and gelatin zymography experiments.

### RT-PCR, western blotting, zymography and Co-IP

Total RNA was isolated using TRIzol (Cat# 15596026, Invitrogen) reagent, followed by cDNA synthesis using High-capacity cDNA synthesis KIT (Cat# 4368814, Thermofisher). Next, the cDNA was used in qPCR using PowerUp^TM^ SYBR^TM^ Green Master Mix (Cat# A25742, Applied Biosystem) and gene-specific primers (Supplementary Table 1) using Quantstudio^TM^ 5. Cyclophilin A (CycA) gene was used as an internal housekeeping gene. Gene fold change was calculated using the ΔΔC_t_ technique [33].

For western blotting, cell lysates were prepared using RIPA buffer (Sigma Aldrich Cat. No. R0278) with 1% protease and phosphatase inhibitors. Equal amounts of denatured proteins were loaded into 10% PA-gels and run for 2 hours at 90V. After protein transfer onto methanol-activated PVDF membranes (Cat# 1620177, Bio-Rad), membranes were blocked using a 1% gelatin solution for 1 hour at RT. The following primary antibodies were used: mouse anti-GAPDH antibody (Cat# AC033, ABclonal), rabbit anti-MMP9 antibody (Cat# MA5-45511, Invitrogen), rabbit anti-ITGβ1 antibody (Cat# PA578028, Invitrogen), and rabbit anti-pMLC antibody (Cat# 3674, Cell Signaling). The following secondary HRP-conjugated antibodies were used: anti-rabbit IgG (Cat. No. 31460, Invitrogen) and HRP-conjugated anti-mouse IgG (Cat. No. 31430, Invitrogen). Finally, the blots were developed in a ChemiDoc^TM^MP Gel Imaging System (Biorad) with ECL substrate solution (Cat. No. K-12045-D20, Advansta). The Fiji-ImageJ software was used to perform densitometric analysis.

Gelatin zymography was performed using cell-secreted conditioned media (CM) to investigate the secretion profile of MMP2/9 by different cell types as described previously [30]. Briefly, the CM collected 24 hours after culture, was run in 10% PA gels containing 0.1% gelatin type B (Cat# 83740, SRL Chemical) using a non-denaturing sample loading buffer. The samples were run at a voltage of 130V and at 4^0^C until the dye front exited the bottom of the gels. After rinsing with renaturing buffer (2.5% TritonX-100 in distilled water) at RT, the gels were incubated in developing buffer (50 mM Tris-base, 50 mM Tris-HCL, 0.2 mM NaCl, 5 mM CaCl2 and pH adjusted to 7.4) for 24 hours at 37^0^C. Next, after incubation for 1 hour at RT with staining solution (40% methanol, 10% acetic acid, 0.1% Coomassie Brilliant Blue in distilled water), the gels were rinsed with detaining solution (40% methanol, 10% acetic acid in distilled water) until the degradation bands became visible. Gel images were captured using Bio rad’s ChemiDoc^TM^MP Gel Imaging System.

The Protein-G CoIP kit (IP50, Sigma Aldrich) was used for coimmunoprecipitation (CoIP) experiments. After lysing cells in non-denaturing lysis buffer (20 mM Tris HCl at pH 8, 137 mM NaCl, 10% glycerol, 1% NP-40, and 2 mM EDTA), cell lysates were incubated overnight with either the anti-ITGβ1 antibody (Cat# PA578028, Invitrogen) or the control IgG antibody (Cat# CR1, Sino Biological Inc.) in an IP spin column at 40 °C. Next, 30µL of Sepharose beads were added and incubated again overnight at 40 °C. After spin, the beads were washed 6-7 times with IP buffer. Finally, the bound protein was eluted by incubating the beads with 1x Laemmli buffer at 95^0^C for 15 minutes. SDS-PAGE was used to load 20 µL of eluted protein and equal amounts of whole cell lysate. Western blotting was performed using antibodies against MMP9 (Cat# MA5-45511, Invitrogen) and GAPDH (Cat# AC033, ABclonal).

### Immunocytochemistry (ICC) & Microscopy

For staining, cells were fixed with 4% paraformaldehyde (PFA) (Cat# 158127, Sigma Aldrich) for 15 minutes at room temperature (RT), washed three times with DPBS, and permeabilized with 0.25% TritonX-100 (Cat# 64518, SRL Chemicals) for 10 minutes at RT. After blocking with 5% BSA (Cat# 1650500501730, Genei) for 1hr at RT, cells were incubated overnight at 40C with one or more of the following antibodies: rabbit anti-pMLC (ser19) (Cat# 3675, Cell Signaling), rabbit anti-paxillin (Cat# PA534910, Invitrogen), rabbit anti-ITGβ1 (Cat# PA578028, Invitrogen), and rabbit anti-MMP9 (Cat# MA5-45511, Invitrogen). In addition, cells were incubated with Wheat Germ Agglutinin (WGA) conjugated with Alexa Flour (AF) 647 (Cat# W32466, Invitrogen) to stain cell glycocalyx and phalloidin conjugated with Alexa Flour (AF) 555 (Cat# A34055, Invitrogen) to stain F-actin stress fibre. The next day, after washing with DPBS, cells were incubated with one of the following secondary antibodies at RT for 2 hours: anti-rabbit Alexa Flour-488 conjugated antibody (Cat# A-11008, Invitrogen), anti-rabbit Alexa Flour-555 conjugated antibody (Cat# A-21428), anti-rabbit Alexa Flour-647 conjugated antibody (Cat# A-31573, Invitrogen), and anti-mouse. The coverslips were treated with DAPI (Cat# D9542, Sigma Aldrich) for 15 minutes at RT after three DPBS washes. After mounting coverslips, samples were imaged using an Olympus IX83 inverted fluorescence microscope and/or an LSM 780 Carl Zeiss confocal microscope at 60x magnification. Images were analyzed with Fiji-ImageJ. Filament-Sensor software [34] was used to measure F-actin stress fibre length and count.

To conduct the Proximity Ligation Assay (PLA) in order to identify the interactions between ITGβ1 and CD44, the Duolink® In Situ Red Starter Kit Mouse/Rabbit (DUO92101, Sigma-Aldrich) was used, following the manufacturer’s instructions. In short, the cells were fixed with 4% PFA for 10 minutes, washed twice with PBS followed by permeabilization with 0.25% Triton-X100 solution for 15 minutes. After washing with PBS twice, the cells were blocked for one hour at 37^0^C in a humid chamber using the supplied blocking solution in the kit. Next, an overnight incubation at 4^0^C was carried out for the primary antibodies against anti-MMP9 and anti-integrin β1. After that, the cells were washed twice using 1x washing solution A. Subsequently, one-hour incubation of the Duolink PLUS/MINUS secondary antibody in a humid chamber at 37^0^C was performed. After washing twice with solution A In a humid 37^0^C chamber, a ligation reaction was carried out for 30 minutes using the Duolink ligation regent. This was followed by an amplification step using Duolink amplification reagents for two hours in the dark humid 37^0^C chamber. Wash solution B was used for the last washing, and Duolink DAPI-containing mounting media was used for the mounting process. Images were captured in an Olympus IX83 inverted fluorescence microscope at 60x magnification.

Focal adhesion (FA) dynamics were assessed by paxillin-mRFP overexpressing cells seeded at low density on glass bottomed dishes (Cat# D35-C4-20-1.5N, CellVis). Time-lapse images were collected using a spinning disc confocal microscope (Yokogawa Electric Corporation, CSU-X1) at 15s intervals for a duration of 10min. After subtracting the background in Fiji-ImageJ, adhesion lifetimes were estimated using the open access Focal Adhesion Analysis Server (FAAS) [35].

For studying the localization dynamics of MMP9 and ITGβ1, GFP-fused MMP9-expressing cells were transfected with ITGβ1-mRFP cDNA. These cells were seeded on glass-bottomed dishes at low density for imaging. Time-lapse images were collected on a spinning disc confocal microscope at 5s intervals for 10 min at low laser power to prevent bleaching. Analysis was performed using Fiji-ImageJ. After subtracting the background, ITGβ1 foci were tracked, and diffusion speed was quantified using manual tracking. Additionally, the ratio of the ITGβ1 foci moving towards the cell migration front to the foci internalized was quantified. For kymograph experiments, cells were sparsely seeded on collagen-coated 24 well plates, and images of live cells were acquired at 5 s intervals for 15 minutes using a phase-contrast microscope (Olympus IX83, 20x objective). Protrusion and retraction rates were quantified using the previously described protocols [36].

### Single cell biophysics measurements

For 2D cell motility experiments, a total of 15,000 cells were seeded onto each 12mm polyacrylamide (PA) gel coverslips. After allowing cells to settle for 12 hours, images were captured at 20-minute intervals over 12 hours using an Olympus IX83 inverted microscope equipped with a 10x objective. For 3D collagen invasion experiments, cells were encapsulated within 1.2 mg/ml 3D collagen matrices as done elsewhere [30], [37]. Timelapse images were recorded at 20 min intervals for 12 hours. This was accomplished using an Olympus IX83 inverted microscope equipped with a 10x objective lens. The velocity of cells in both 2D motility and 3D invasion was measured using the manual tracking plug-in of the Fiji-ImageJ software. Subsequently, an Excel macro, as detailed in a separate publication [38], was utilized.

For gel compaction assay, 5x10^4^ cells were copolymerized with 1.2mg/mL collagen (Cat# 354236, Corning) to a total volume of 150µL in 48 well plates. After 24hr, the polymerized gels were released from the sides using a needle. After 48hr, gel images were captured in an Olympus IX83 phase contrast microscope at 4x magnification. The quantification of the gel area was done in Fiji-ImageJ software using the following equation:

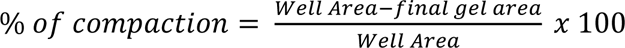

Atomic force microscopy (AFM) was used to measure the cortical stiffness of cells. Pyramidal AFM probes (10 kHz, Cat # TR400PB, Olympus) with a nominal spring constant of 30pN/nm were calibrated thermally to obtain the precise value of the spring constant. Cells were grown at low density on stiff PA gels (5 kPa) on 12mm coverslips for 12-16 hours. Indentation was done in the cell cortex area to measure the force curve, and the data was fitted in the Hertz model till 500nm.

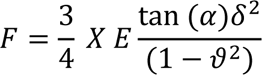

Here, F = indentation force, E = elastic modulus of the material, α = half tip angle, δ = indentation depth, ν = Poisson ratio of the material (assumed to be 0.45 for the cell).

Adhesion tests using AFM were carried out to detect RGD-ITGβ1 interactions. Cells were grown on 5 kPa stiff PA-gels for 12-16h. The spherical probe (67 kHz, Novascan) was coated with 10µg/mL poly-L-lysin solution for 12 hours and treated with 0.5% glutaraldehyde for 4 hours. Ultrapure water is used three times to wash the probes to remove surplus solutions. Next, the probe was treated with 10µg/mL RGD peptide for 2 hours. The probe was rinsed with ultrapure water to eliminate unbound RGD. During the experiment, the functionalized probe was held on the cell surface for 10s to generate RGD integrin bonds, then retracted at 4µm/s. In this step, we recorded the retraction curve and estimated the adhesion force using Igor Pro and MATLAB.

### Statistical Analysis

The Kolmogorov-Smirnov normality test was used to examine data distribution. A parametric or nonparametric statistical test was done based on the outcome. For parametric data, one-way ANOVA was employed for statistical analysis, and Tukey’s test was utilized to compare means. For non-parametric data, the Mann-Whitney test was used. Prism 9 (GraphPad Software, LLC) was used for statistical analysis and generating graphs, with P<0.05 considered statistically significant.

## Results

### ECM stiffness differentially modulates MMP expression and activity in cancer cells

Pan-cancer analysis of the TCGA database using the UCSC Xena browser (Fig. 1A) revealed elevated expression of MMP1, 9, and 14 in primary and secondary tumor sites compared to solid benign tumors, with MMP2 transcript levels remaining unchanged (Fig. 1B). Similar trend observed in breast and lung cancers establishes a strong positive correlation between overexpression of MMP1, 9, and 14 with tumorigenicity and malignancy (Fig. 1C, D).

**Figure 1:**
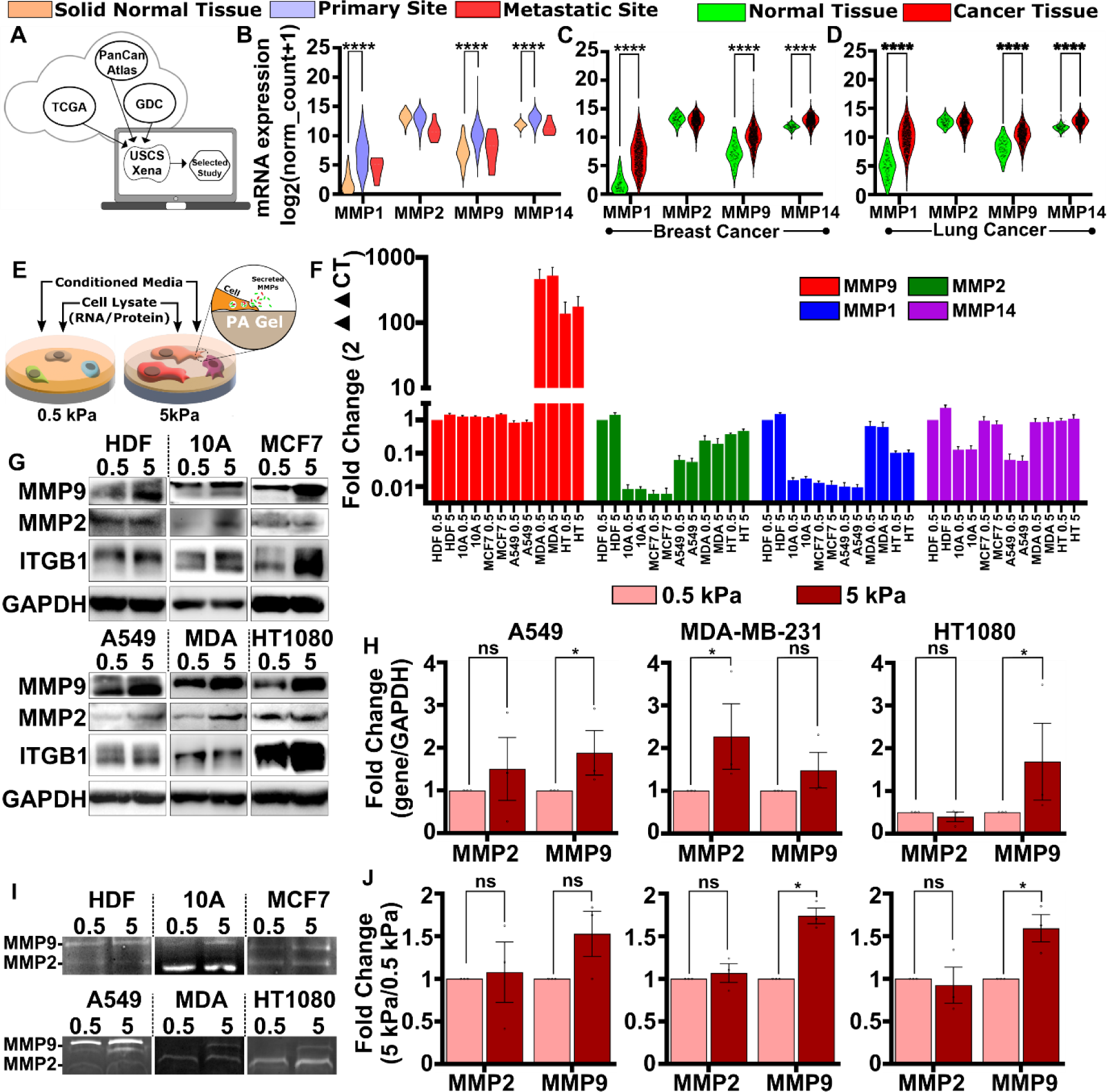
ECM stiffness differentially modulates MMP expression and activity in cancer cells. (A) Curing TCGA patient datasets and processing to assess the expression profile of major MMPs (MMP1, 2, 9, and 14) using UCSC Xena browser. (B) The mRNA expression profiles of MMP1, 2, 9, and 14 in ∼110 normal tissue samples, ∼1100 tissues from primary malignant sites, and ∼8 tissues from secondary malignant sites. (C, D) The expression profiles of MMP1, 2, 9, and 14 in ∼115 normal and ∼1100 malignant tissue samples in breast cancer patients and lung cancer patients. (E) Schematic for assessing stiffness dependent MMP expression and secretion. Cells were grown on 0.5 kPa soft and 5 kPa stiff polyacrylamide gels. (F) Relative fold change of gene expression of MMP1, 2, 9, and 14 by quantitative PCR in human dermal fibroblasts (HDFs), MCF10A normal mammary epithelial cells, MCF7 ERPR+ breast cancer cells, A549 lung adhenocarcinoma cells, MDA-MB-231 (MDA) triple negative breast cancer cells, and HT1080 (HT) fibrosarcoma cells on both 0.5 and 5 kPa PA-gels. The fold change was calculated for the expression of HDFs on 0.5 kPa gels. (G) Western blot analysis of protein expression of MMP2, 9, and ITGβ1 in the cells mentioned above on 0.5 and 5 kPa PA-gels. GAPDH was used as a loading control. (H) The fold change of the gene expression of A549, MDA-MB-231, and HT1080 cells on 5 kPa PA gels normalized with GAPDH expression is represented as a bar diagram analyzed from three independent experiments (*mean* ± *SEM*). (I) Gelatin zymogram of cell-secreted conditioned media (CM) from soft and stiff substrates to probe the secretion profiles of MMP9 and MMP2. (J) The relative fold change of the secreted MMP2/9 of A549, MDA-MB-231, and HT1080 on 5 kPa PA gels is represented as the bar diagram normalized to on 0.5 kPa PA gels. The analysis was performed three independent times and densitometric analysis of the band intensity was calculated for statistical comparisons (*mean* ± *SEM*). For all sub-figures, statistical significance was assessed using One-Way ANOVA or nonparametric test. (∗ *p* < 0.05, *ns*: *not significant*).

Since matrix stiffening is implicated in cancer progression [30], stiffness dependent expression profiling of MMP1/MMP2/MMP9/MMP14 was carried out on soft (0.5 kPa) and stiff (5 kPa) polyacrylamide (PA) gels mimicking normal and metastatic tissue stiffness (Fig. 1F) [30]. Experiments were performed using normal human dermal fibroblasts (HDFs), non-cancerous MCF10A breast epithelial cells, non-invasive ER^+^PR^+^ MCF7 breast cancer cells, moderately invasive A549 lung adenocarcinoma cells, and highly invasive MDA-MB-231 triple-negative breast cancer (TNBC) cells and HT1080 fibrosarcoma cells (hereafter referred to as MDA and HT cells, respectively). While MMP1, MMP2, and MMP14 levels were comparable across HDFs, MDA, and HT cells, more than 100-fold MMP9 overexpression was observed in MDA and HT cells (Fig. 1F). In addition, MDA and HT cells also exhibited ∼2-fold stiffness dependent MMP9 upregulation on 5 kPa gels compared to 0.5 kPa gels. Consistent with these observations, the western blot of cell lysates revealed elevated expression of MMP9 and the mesenchymal marker integrin-β1 in all the cells cultured on 5 kPa gels (Fig. 1G, H and Supp. Fig. 1A). Stiffness-mediated MMP2 upregulation was observed in MCF-10A and MDA only. Assaying MMP2/MMP9 secretion using cell-secreted conditioned media (CM), revealed significant stiffness dependent MMP9 activity in gelatin zymogram from highly invasive MDA and HT cells only, with no stiffness dependent alteration in MMP2 activity (Fig. 1I, J and Supp. Fig. 1B). Collectively, these results suggest that high baseline MMP9 expression and stiffness dependent MMP9 modulation are hallmarks of highly metastatic cancer cells.

### Soluble MMPs drive cell spreading and migration by regulating focal adhesions

To probe the role of soluble MMPs (i.e., MMP2 and MMP9) in mediating stiffness dependent invasive phenotype, we measured cell spreading and motility of A549, MDA, and HT cells on soft (0.5 kPa) and stiff (5 kPa) PA gels in the presence and absence of the pan-MMP inhibitor GM6001 (hereafter referred to as GM) and the soluble MMP2/9 inhibitor SB3-CT (hereafter referred to as SB3). DMSO-treated and untreated cells served as controls. While both GM and SB3-treated cells exhibited a dramatic drop in cell spreading and increased cell rounding on stiff PA gels (Fig. 2A, B and Supp. Fig. 2C), no significant changes were observed on soft PA gels (Supp. Fig. 2A, B and C). In line with these observations, cell motility was markedly reduced on stiff gels (Fig. 2C, D, Supp. Fig. 2E and Supp. Movie 2) with no changes on the soft gels (Supp. Figs. 2D, F and Supp. Movie 1). Similarly, when encapsulated in 3D collagen gels, cell invasiveness was inhibited in GM and SB3 treated cells (Fig. 2E, F, Supp. Fig. 2G, and Supp. Movie 3).

**Figure 2:**
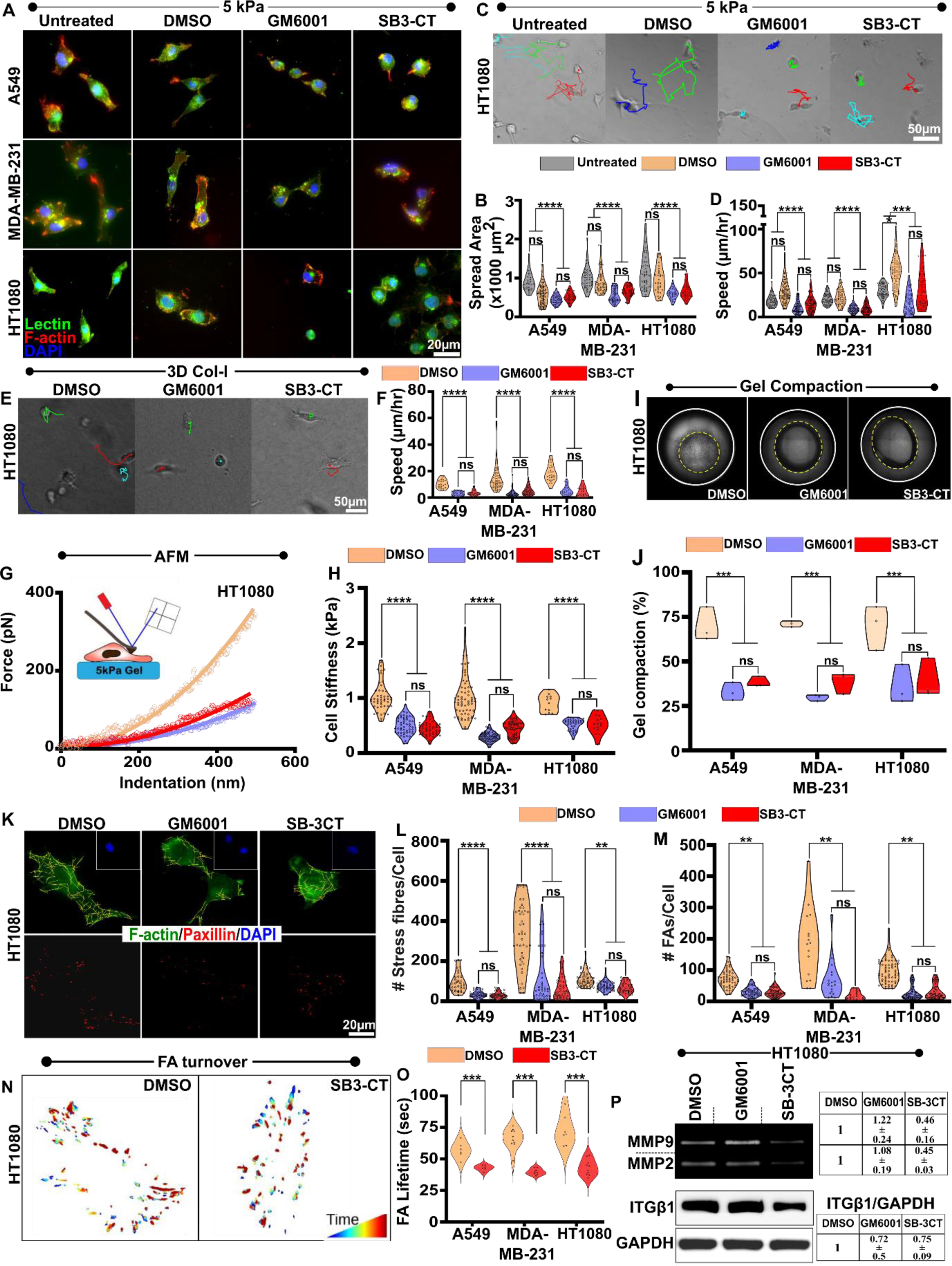
Soluble MMPs drive cell spreading and migration by regulating focal adhesions. (A, B) Representative Lectin (green)/F-actin (red)/DAPI (blue) stained images of untreated and drug treated HT1080 cells on 5 kPa PA gels and quantification of cell spreading area of untreated, control (DMSO) and drug (GM6001 and SB3-CT) treated A549, MDA-MB-231 and HT1080 cells (≈ 80 − 100 cells, *N* = 3). (C, D) Representative migration trajectories of control and drug treated HT1080 cells on 5 kPa PA gels, and quantification of cell speed (≈ 50 − 80 cells per condition, *N* = 3). (E, F) Representative invasion trajectories of control and drug treated HT1080 cells entrapped inside 3D collagen gels and quantification of cell speed (≈ 30 − 50 cells per condition, *N* = 3). (G, H) Representative force vs. indentation curves of control and drug treated HT1080 cells on 5 kPa PA gel. The data were fitted to obtain estimates of cell cortical stiffness (≈ 40 − 50 cells per condition, *N* = 3). (I, J) Representative images of HT1080 cells/Collagen I gel mixture cultured for 48 hr in the presence of DMSO/GM6001/SB3-CT and quantification of gel compaction (≈ 3 − 4 gels per condition, *N* = 3). White/yellow dotted circles represent the initial/final gel areas. (K-M) Representative images of F-actin stress fibre (green) and paxillin (red) stained HT1080 cells treated with DMSO, GM6001 and SB3-CT on 5 kPa PA gels, and quantification of stress fibers and focal adhesion size (≈ 30 − 50 cells per condition, *N* = 3). The yellow lines represent processed stress fibres. (N, O) Representative lifetime heatmaps and quantification of focal adhesion lifetime of HT1080 cells treated with DMSO and SB3-CT (≈ 7 − 10 cells per condition, *N* = 3). (P) The gelatin zymogram of conditioned media (CM) and western blot of ITGβ1 in DMSO, GM6001 and SB3-CT treated HT1080 cells. Densitometric analysis of the bands were performed (*N* = 3). GAPDH was used as a loading control. For all sub-figures, statistical significance was assessed using One-Way ANOVA or nonparametric test. (∗ *p* < 0.05,∗∗ *p* < 0.01,∗∗∗ *p* < 0.001,∗∗∗∗ *p* < 0.0001, *ns*: not significant.

Defects in migration can be attributed to reduced cytoskeletal forces and/or perturbed adhesion formation and dynamics. Both GM and SB3 treatment led to prominent cell softening in A549, MDA, and HT cells (Fig. 2G, H and Supp. Fig. 2I), and ∼50% drop in the extent of gel compaction (Fig. 2I, J and Supp. Fig. 2H) indicative of a compromised cytoskeleton, as evident from a marked reduction in the number of F-actin stress fibres/cell (Fig. 2K, L and Supp. Fig. 2J). Since focal adhesions stabilize the cytoskeleton, we next probed adhesion size distribution and adhesion stability in GM/SB3 treated cells. Visualization of paxillin-stained focal adhesions revealed a dramatic drop in the average number of focal adhesions in drug-treated cells (Fig. 2K, M and Supp. Fig. 2J). Furthermore, time-lapse images of cells stably transfected with GFP-paxillin revealed a marked reduction in focal adhesion (FA) lifetime upon SB3 treatment (Fig. 2N, O, Supp. Fig. 2K and Supp. Movie 4). Reduction in FA stability was associated with a drop in the secretion of MMP2 and MMP9 and a reduction in ITGβ1 expression (Fig. 2P) in HT1080 cells. Together, these results suggest that cell secreted soluble MMPs mediate cell spreading and migration by modulating adhesion stability and turnover.

### Both proteolytic and non-proteolytic functions of MMP9 are essential for sustaining cancer invasion

In contrast to SB3, treating MDA and HT cells with ARP100 at concentrations that inhibited only MMP2 [31], did not impact cell spreading and motility (Supp. Fig. 2L, M and Supp. Movie 5), suggesting that defects in cell spreading and motility observed in SB3-treated cells is attributed to MMP9. To test the role of MMP9 directly, we established shCTL (i.e., cells transfected with scrambled shRNA) and shKD1/shKD2 knockdown cells (i.e., MMP9 was knocked down using two distinct shRNAs) in HT cells. Knockdown cells were re-transfected with MMP9 full-length cDNA to generate M9FL1 and M9FL2 cells for recovery experiments. For probing non-proteolytic functions of MMP9, knockdown cells were also transfected with catalytically inactive MMP9 (E402A) cDNA to generate ΔCAT cells (Fig. 3A, Supp. Fig. 3A and B) [39]. While comparable levels of MMP9 were detected in conditioned media (CM) secreted by M9FL1/M9FL2 cells, degradation bands in gelatin zymograms were detected in shCTL and M9FL cells only (Fig. 3B).

**Figure 3:**
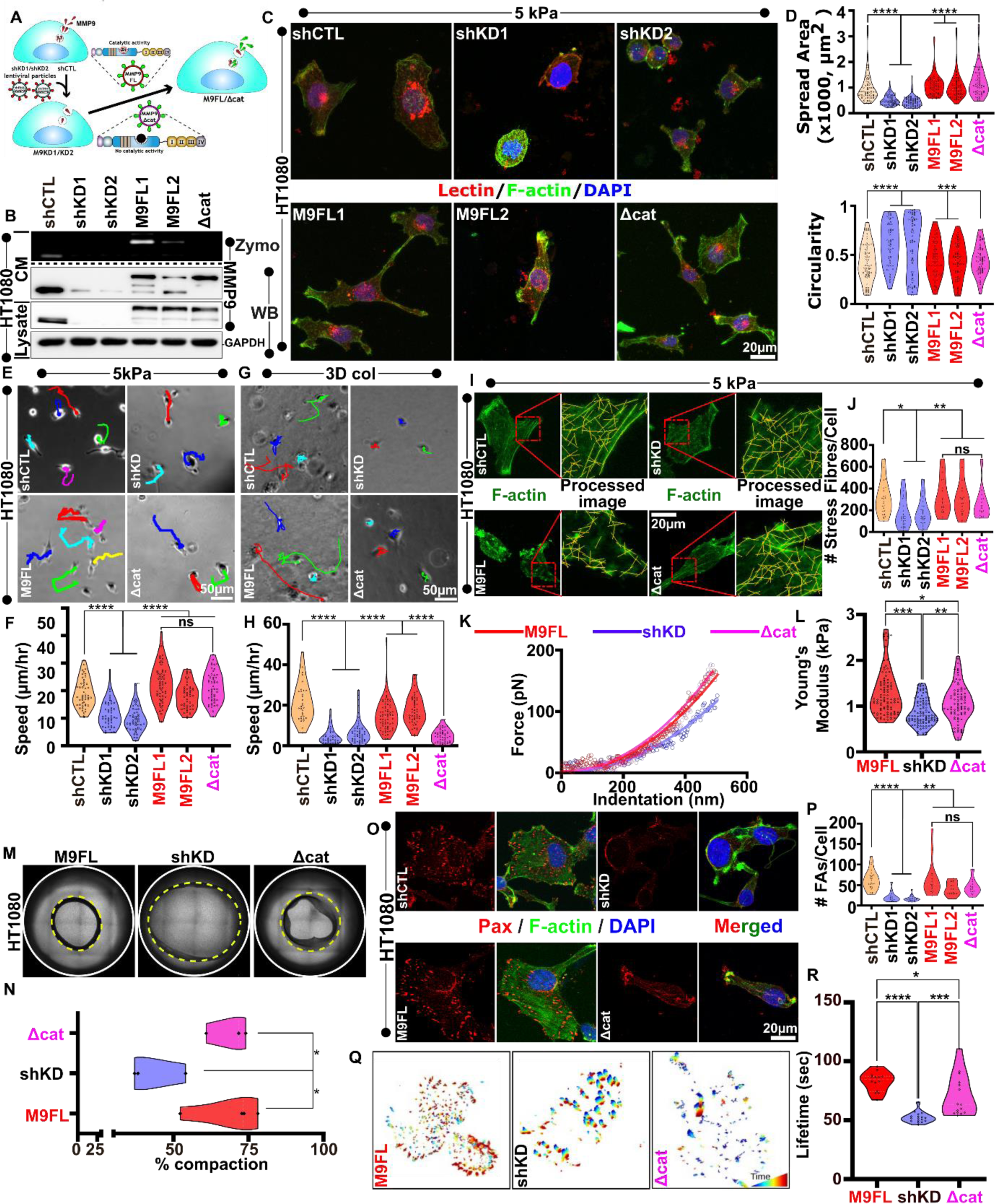
Both proteolytic and non-proteolytic functions of MMP9 are essential for sustaining cancer invasion. (A) Schematic of generation of MMP9 knockdown cells (shKD1/2) and M9FL/Δcat expressing cell lines using lentiviral mediated shRNA transduction. (B) Gelatin zymogram of CM and western blotting of CM/ whole cell lysate from shCTL, shKD1/2, M9FL1/2 and Δcat cells. (C, D) Representative lectin (green), phalloidin (red) and DAPI (blue) stained images of HT1080 shCTL, shKD1/2, M9FL1/2 and Δcat images of cells on stiff 5 kPa PA gels, and quantification of cell spread area and circularity (≈ 50 − 80 cells per condition, *N* = 3). (E, F) Representative 2D migration trajectories of HT1080 shCTL, shKD, M9FL and Δcat cells on 5 kPa PA gels and quantification of cell motility (≈ 50 − 80 cells per condition, *N* = 3). (G, H) 3D invasion trajectories of HT1080 shCTL, shKD, M9FL and Δcat cells inside 3D Col and quantification of cell speed (≈ 40 − 50 cells per condition, *N* = 3). (I, J) Representative images of F-actin (green) of HT1080 shCTL, shKD, M9FL and Δcat cells, and quantification of stress fibers (≈ 50 − 60 cells per condition, *N* = 3). Yellow lines indicate individual stress fibres. (K, L) Representative force-indentation of M9FL, shKD and Δcat cells and quantification of the cortical stiffness of these cells (≈ 50 − 80 cells per condition, *N* = 3). (M, N) Representative images of 3D gel compaction of M9FL, shKD and Δcat cells and quantification of gel compaction (≈ 3 − 4 gels per condition from *N* = 3). White/yellow dotted circles represent the initial/final gel areas. (O, P) Representative paxillin (red) and F-actin (green) stained shCTL, shKD, M9FL and Δcat cells and quantification of the number of focal adhesions per cell (≈ 30 − 50 cells per condition, *N* = 3). (Q, R) Representative Focal adhesion lifetime heatmaps and quantification of adhesion lifetime in M9FL, shKD and Δcat cells (≈ 15 − 20 cells per condition, *N* = 3). For all sub-figures, statistical significance was assessed using One-Way ANOVA or nonparametric test (∗ *p* < 0.05,∗∗ *p* < 0.01,∗∗∗ *p* < 0.001,∗∗∗∗ *p* < 0.0001, *ns*: not significant).

In contrast to shCTL cells which were well spread and polarized, shKD1/shKD2 cells exhibited reduced spreading with rounded morphologies similar to SB3 treated cells (Fig. 3C and D). Remarkably, cell spreading was rescued not only in M9FL1/M9FL2 cells but also in ΔCAT cells. In line with these observations, reduction in cell motility on 5 kPa gels observed in shKD1/shKD2 cells, was rescued in both M9FL1/M9FL2 and ΔCAT cells (Fig. 3E, F and Supp. Movie 6). However, when entrapped in 3D collagen gels, in contrast to shCTL and M9FL cells which migrated at comparable speeds, both shKD1/shKD2 and ΔCAT cells were unable to migrate (Fig. 3G, H and Supp. Movie 7). These results highlight the importance of MMP9 catalytic activity in sustaining invasion through dense matrices.

To probe the non-proteolytic functions of MMP9, cytoskeletal organization and focal adhesion dynamics were compared across the different conditions. Reduction in stress fibre formation in shKD1/shKD2 cells (Fig. 3I, J) was associated with reduced actomyosin contractility (assessed using pMLC staining) (Supp. Fig. 3C and D) and gel compaction (Fig. 3M and N), and increased cell softening (Fig. 3K, L). However, re-expression of full length MMP9 or MMP9 ΔCAT restored cytoskeletal organization, contractility, and cell stiffness to levels comparable to shCTL cells.

Cell migration involves cycles of protrusion and adhesion stabilization at the cell front. Since kymograph analysis revealed perturbed protrusion dynamics in shKD1/shKD2 cells only, but not in ΔCAT cells (Supp. Fig. 3E-G), we hypothesized MMP9 ΔCAT is fully capable of stabilizing adhesions. Indeed, in direct contrast to shKD1/shKD2 cells where the frequency of focal adhesions and their stability were significantly reduced, both focal adhesion number and lifetime were comparable in M9FL and ΔCAT cells (Fig. 3O-R and Supp. Movie 8). Together, these results suggest that proteolytic and non-proteolytic functions of MMP9 collectively sustain cancer invasion, with MMP9 stabilizing adhesions in a non-proteolytic manner and proteolytic remodeling of the matrix.

### MMP9-integrin **β1** (**ITGβ1**) interaction via two distinct sites is essential for sustaining cancer invasion

How does MMP9 stabilize adhesions? In a previous study, hemopexin domains of MMP14 have been shown to mediate binding with integrin β1 (ITGβ1) and activate MAPK [40]. Another study identified a 17-amino acid motif in blade IV of the hemopexin domain that mediates integrin binding [29] (Fig. 4A). MMP9 catalytic domain is also known to harbour three fibronectin type II like repeats [41], [42]. Bioinformatics analysis revealed the presence of only one RGD motif in the catalytic domain of MMP9 (Fig. 4A). Based on the background literature, and our bioinformatics analysis, we speculated MMP9 might regulate focal adhesions non-proteolytically through its interactions with ITGβ1 (Fig. 4A). To test our hypothesis, we generated two mutations at these sites: an RGD-to-RGE substitution and a deletion of 17 amino acids in the PEX4 domain (Δβ1) (Fig. 4B). This was done in both M9FL and ΔCAT cells, with these cell lines referred to as RGE, Δβ1 and Δβ1ΔCAT cells, respectively. Thus, while matrix proteolysis is expected to remain unchanged in M9FL, RGE and Δβ1 cells, integrin binding is expected to be perturbed in RGE, Δβ1 and Δβ1ΔCAT cells (Fig. 4B). Co-immunoprecipitation (CoIP) experiments confirmed that interaction between MMP9 and ITGβ1 was eliminated in Δβ1 and Δβ1ΔCAT cells, but not in M9FL, ΔCAT and RGE cells (Fig. 4C).

**Figure 4:**
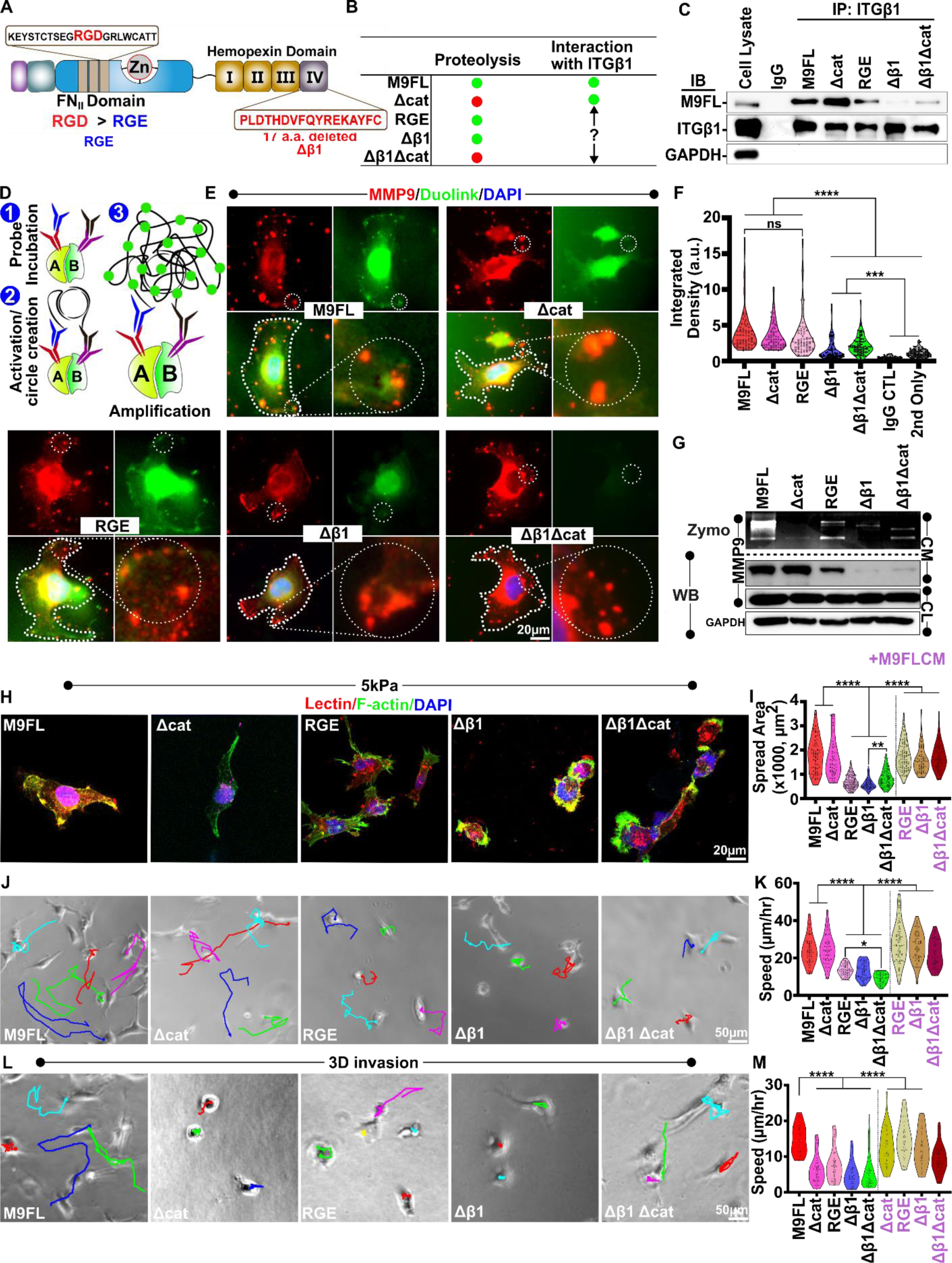
MMP9-ITGβ1 interaction via two distinct sites is essential for sustaining cancer invasion. (A) Domain organization of MMP9 showing two putative integrin bindings sites, the RGD sequence in the catalytic domain and a 17 aa sequence in blade IV of the hemopexin domain. To assess the importance of these two sites, two mutants RGE and Δβ1 were generated. (B) Proposed proteolytic and nonproteolytic functions of different mutated MMP9 constructs, where green/red circles represent active/inactive functions. (C) Co-immunoprecipitation for the verification of the perturbation of association between different MMP9 constructs (M9FL, Δcat, RGE, Δβ1, and Δβ1Δcat) and ITGβ1. (D) The working principle of proximity ligation assay (PLA) using Duolink Probe. (E, F) Spatial localization of MMP9-ITGβ1 association determined by PLA (green) and MMP9 (red) signal in M9FL, Δcat, RGE, Δβ1, and Δβ1Δcat cells and quantification of PLA integrated density (≈ 60 − 80 cells per condition, *N* = 3). IgG control and only secondary antibody-treated conditions served as negative controls. (G) MMP9 secretion and expression in M9FL, Δcat, RGE, Δβ1, and Δβ1Δcat cells assessed using gelatin zymography and western blotting of CM/whole cell lysate. GAPDH served as loading control. (H, I) Representative lectin (red)/F-actin (green)/DAPI (blue) stained images of M9FL, Δcat, RGE, Δβ1, and Δβ1Δcat cells and quantification of cell spread area (≈ 70 − 80 cells per condition, *N* = 3). (J-M) Representative 2D migration and 3D invasion trajectories of M9FL, Δcat, RGE, Δβ1, and Δβ1Δcat cells on 5 kPa PA gel and 3D collagen respectively and quantification of cell speed (≈ 60 − 80 cells per condition, *N* = 3). For all sub-figures, statistical significance was assessed using One-Way ANOVA or nonparametric test (∗ *p* < 0.05,∗∗ *p* < 0.01,∗∗∗ *p* < 0.001,∗∗∗∗ *p* < 0.0001).

Proximity ligation assay (PLA) enabled us to visualize how MMP9/ITGβ1 association was spatially distributed (Fig. 4D). While co-localization of MMP9 (red) with PLA signal (green) was prominently observed in M9FL, ΔCAT and RGE cells at the cell periphery (marked by white dotted circles), this was markedly reduced and/or absent in Δβ1 and Δβ1ΔCAT cells, as evident from quantification of the integrated density of the PLA signal (Fig. 4E, F).

To next assess if MMP9/ ITGβ1 association influenced MMP9 secretion, gelatin zymography experiments were performed using cell secreted conditioned media (CM) from the five cell lines. As expected, MMP9 band was prominently detected in M9FL cells only (Fig. 4G). While the absence of MMP9 band in ΔCAT cells was expected, MMP9 band intensities were substantially reduced in both RGE and Δβ1 cells. Consistent with this, no MMP9 bands were detected in western blotting of CM secreted by RGE, Δβ1 and Δβ1ΔCAT cells, suggesting that MMP9/ ITGβ1 interaction is required for MMP9 secretion.

To assess the functional importance of MMP9/ ITGβ1 association, cell spreading, motility and invasion experiments were performed as described above. Loss of MMP9/ ITGβ1 association in RGE, Δβ1 and Δβ1Δcat cells led to pronounced reduction in cell spreading and increase in cell rounding (Fig. 4H, I, Supp. Fig. 4A). However, the addition of CM from M9FL cells led to near complete recovery in cell spreading and polarized morphology. RGE, Δβ1 and Δβ1Δcat cells, which exhibited minimal spreading, also exhibited defective motility, which was rescued upon the addition of M9FL CM (Fig. 4J, K, Supp. Fig. 4C and Supp. Movie 9 and 10). Similar results were obtained when cells were entrapped in 3D collagen gels and their invasiveness was tracked, i.e., RGE, Δβ1 and Δβ1Δcat cells were unable to invade along with Δcat, but invasiveness was rescued by adding M9FL CM (Fig. 4L, M, Supp. Fig. 4D and Supp. Movie 11 and 12). Taken together, these results suggest that MMP9/ ITGβ1 association is essential for sustaining cancer invasion.

### MMP9-integrin association via PEX4 domain mediates ITGβ1 membrane trafficking while RGD domain of MMP9 mediates ITGβ1 membrane stabilization

To understand the mechanisms underlying defective spreading and motility of RGE, Δβ1, and Δβ1Δcat cells, we first assessed if adhesion formation/dynamics and/or cytoskeletal organization were negatively impacted. Paxillin and stress fiber staining showed that disruption of MMP9/ITGβ1 association led to a reduction in both the number focal adhesions as well as the number of stress fibers in RGE, Δβ1, and Δβ1Δcat cells (Fig. 5A, B, Supp. Fig. 5A and B). Timelapse imaging of paxillin-eGFP transfected cells revealed adhesion stability was reduced in RGE, Δβ1 and Δβ1Δcat cells compared to M9FL/Δcat cells (Fig. 5C, D, Supp. Movie 13).

**Figure 5:**
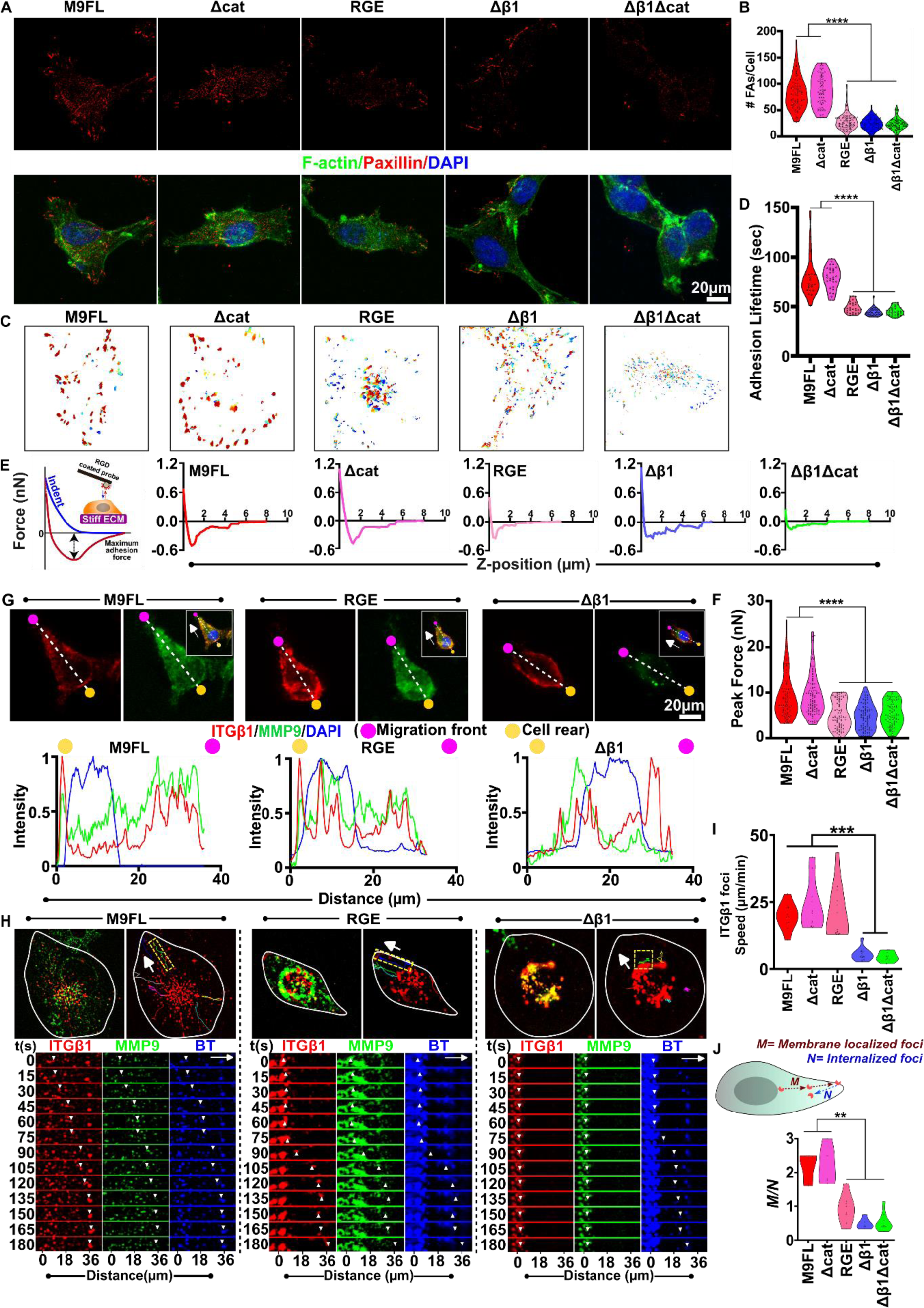
ITGβ1 binding sites on MMP9 play distinct roles. (A, B) Representative paxillin (red)/F-actin (green) stained images of M9FL, Δcat, RGE, Δβ1, and Δβ1Δcat cells on 5 kPa PA gels and quantification of number of FAs per cell (≈ 50 − 80 cells per condition, *N* = 3). (C, D) Focal adhesion lifetime heatmaps of M9FL, Δcat, RGE, Δβ1, and Δβ1Δcat cells and its quantification (≈ 12 − 15 cells per condition, *N* = 3). (E, F) Representative AFM retraction curves of M9FL, Δcat, RGE, Δβ1, and Δβ1Δcat cells obtained by indenting cells with RGD-coated spherical tips, and retracting after 10s of dwell time. The maximum adhesion force was quantified for the different cell lines (≈ 50 − 80 cells per condition, *N* = 3). (G) Representative images of ITGβ1 (red), MMP9 (green), and DAPI (blue) stained M9FL, RGE and Δβ1 cells, with intensity along the Line profiles drawn in the direction of cell migration (magenta and yellow circles depict the migration front and cell rear respectively). (H) Trajectories of ITGβ1 (red) foci in M9FL, RGE and Δβ1 cells along the direction of migration (white arrow). Merged images of MMP9 (green) and ITGβ1 foci (red) are also shown. Time stamp images of MMP9 (green), ITGβ1 foci (red) and bio-tracker (blue) of the indicated dotted box (inset) are represented. (I) Quantification of trafficking speed of ITGβ1 foci in the different cell lines (≈ 20 − 30 taken from 5 − 7 cells per condition, *N* = 3). (J) Quantification of the ratio of the number of the ITGβ1 foci going towards the membrane (*M*) to the number of foci being internalized (*N*) at a given time (≈ 15 − 30 taken from 5 − 6 cells per condition, *N* = 3). For all sub-figures, statistical significance was assessed using One-Way ANOVA or nonparametric test (∗ *p* < 0.05,∗∗ *p* < 0.01,∗∗∗ *p* < 0.001,∗∗∗∗ *p* < 0.0001).

Smaller adhesion formation and reduced adhesion stability can be attributed to reduced availability of integrins at the cell membrane and/or their membrane stability. To test this possibility, we performed adhesion experiments using AFM as detailed elsewhere [30]. In brief, cells were indented with RGD functionalized spherical AFM tips; upon indentation, the tip position was held fixed for 10 seconds to allow for adhesions to form, and then retracted (Fig. 5E). Bonds of adhesion formed between the cell surface and the tip, if any, had to be broken for the tip to retract, leading to a characteristic peak in the retraction curve and corresponds to the force required to break the adhesions. As evident from the representative retraction curves (Fig. 5E), compared to M9FL/Δcat cells, the maximum adhesion force was reduced by nearly 50% in RGE, Δβ1 and Δβ1Δcat cells (Fig. 5F), indicative of reduced ITGβ1 localization and/or stability at the cell membrane. Consistent with these observations, MMP9/ITGβ1 dual staining revealed MMP9/ITGβ1 co-localization at the migration front in M9FL and Δcat cells (Fig. 5G, Supp. Fig. 5C). In comparison, MMP9/ITGβ1 colocalization at the cell periphery was reduced in RGE cells, and completely lost in Δβ1/Δβ1Δcat cells, where both MMP9 and ITG β1 exhibited cytoplasmic localization.

Both MMP9 and ITGβ1 are trafficked to the cell membrane via vesicles [43], [44], [45]. Since MMP9 secretion was significantly reduced in Δβ1/Δβ1Δcat cells, we hypothesized that altered localization of both MMP9 and ITGβ1 in Δβ1/Δβ1Δcat cells is attributed to defective vesicular trafficking, with MMP9 and ITGβ1 association ensuring the are being co-transported. To test this, we performed three colour timelapse imaging wherein M9FL, Δcat, RGE, Δβ1 and Δβ1Δcat cells (tagged in green) were transfected with ITGβ1 mRFP, and vesicles were tagged using bio-tracker dye (Blue) (Fig. 5H and Supp. Fig. 5D). Indeed, in M9FL, Δcat and RGE cells, movement of co-localized dots towards the cell periphery is indicative of co-transport of MMP9 and ITGβ1 in the same vesicles (Supp. Movie. 14-18). In contrast, in Δβ1 and Δβ1Δcat cells, co-transport was near completely abolished. Quantification of speed of individual ITGβ1 foci revealed comparable rates of transport in M9FL, Δcat and RGE cells, and substantially reduced transport in Δβ1 and Δβ1Δcat cells (Fig. 5I).

While AFM experiments revealed drop in peak forces in RGE cells, no differences in ITGβ1 transport was observed in these cells. To test if this apparent anomaly can be explained by altered ITGβ1 stability, we tracked the ratio (*M*/*N*) of the number of foci going towards the membrane (*M*) to those being internalized (*N*). While this ratio was around 2 for both M9FL and Δcat cells, this reduced to ∼1 for RGE cells (Fig. 5J), suggesting that in spite of being transported to the membrane, ITGβ1 is not stabilized at the cell membrane in RGE cells. Collectively, these results highlight two distinct roles for the two ITGβ1 binding sites on MMP9, with the 17 amino acid residue essential for ITGβ1 transport and the RGD domain essential for stabilizing ITGβ1 at the cell membrane.

## Discussion

MMP9 expression is associated with increased cancer cell invasion and metastasis [46]. While the proteolytic activity of MMP9 involved in matrix remodeling is well established, an increasing body of literature has demonstrated its non-proteolytic functions play an equally crucial role. Here, we have established how the combination of proteolytic and non-proteolytic functions of MMP9 together render cancer invasiveness via its association with ITGβ1. Based on our findings, we propose a mechanism whereby MMP9/ITGβ1 association is essential for their co-transport to the leading edge, ITGβ1 stabilization at the cell membrane, MMP9 secretion and MMP9 mediated matrix degradation thereby creating migration paths (Fig. 6).

**Figure 6:**
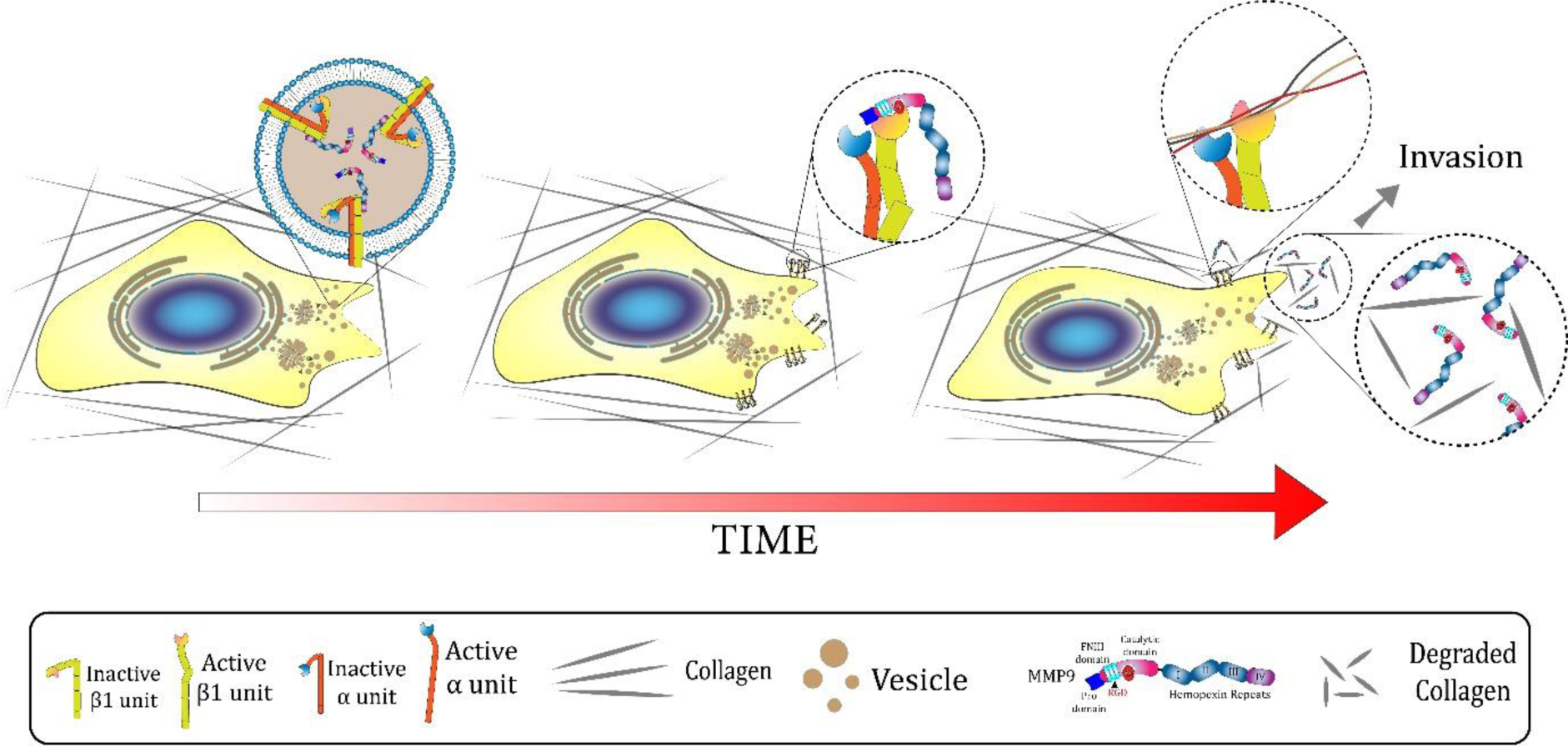
Proposed model of MMP9 mediated ITGβ1 co-trafficking and stabilization on the cell membrane. Both MMP9 and ITGβ1 are co-trafficked via intracellular vesicles to the cell membrane. While hemopexin domain of MMP9 is responsible for co-packaging of MMP9/ITGβ1 in vesicles and their co-transport, RGD domain mediates membrane surface stabilization of ITGβ1. Upon MMP9 mediated matrix degradation, focal adhesions are stabilized leading to forward migration.

The stiffness of the extracellular matrix has a significant impact on cell spreading, motility, cytoskeletal organization, and MMP secretion [30], [47], [48], [49]. Among the secreted MMPs, MMP2 and MMP9, increased cancer invasiveness was associated with a stiffness dependent increase in MMP9 secretion, with MMP inhibition inducing cell rounding, and cell and nuclear softening [50], [51]. By screening both normal and cancer cells of varying invasiveness, here we show that stiffness modulates MMP9 expression transcriptionally in A549, MDA-MB-231 and HT-1080 invasive cancer cells only. However, even in these invasive cells, stiffness dependent modulation of MMP1, MMP2 and MMP14 was not observed. Our results suggest that MMP9 mediates faster motility on stiffer substrates by stabilising adhesions as both pharmacological inhibition of MMPs as well as MMP9 knockdown led to reduction in adhesion size and stability, and destabilization of the cell cytoskeleton [52], [53], [54].

Previous literature has shown that interaction between the MMP9 PEX (PEX9) domain and ITGβ1 is essential for formation of cell-matrix adhesions [26], [27], [55]. Building on this work, another study showed that a 17 amino acid stretch from the fourth blade of the PEX9 domain stabilizes MMP9/ITGβ1 association leading to loss of adhesions. Consistent with these studies, by deleting this stretch of amino acids, we show here that reduced spreading, migration, and invasion of Δβ1 cells is attributed to loss of focal adhesion formation. While the presence of fibronectin type II repeats in MMP9 has been documented before, their role has not been elucidated. For the first time, we identify the presence of an RGD motif in these repeats, which when mutated to RGE, leads to loss of adhesions. Thus, our study establishes that both integrin binding sites on MMP9 are essential for adhesion formation. This MMP9/ITGβ1 association can explain the restoration of the cell spreading, adhesion, and 2D motility in catalytic inactive MMP9 (Δcat) on 2D substrates. But the defects in cell invasion of Δcat cells inside 3D collagen illustrates the necessity of matrix degradation in 3D invasion.

Vesicular transport of MMP9 towards lamellipodia has been reported in HMEC-1 cell invasion previously [56]. Here we show that the 17 amino acid sequence mediates co-trafficking of MMP9 and ITGβ1 to the migration front. Co-trafficking of MMP9 and ITGβ1 ensures that MMPs are secreted at the vicinity of existing adhesions thereby leading to localized matrix degradation and adhesion strengthening. Defective MMP9 secretion in Δβ1 and Δβ1Δcat cells suggests that adhesion maturation and matrix remodeling are closely coupled, with MMPs being released only after integrins are stabilized on the membrane. Reduced MMP9 secretion observed in SB3-CT cells may be attributed to disruption of MMP9/ITGβ1 association. RGD peptide is widely used for integrin blocking experiments. The combination of AFM experiments and time-lapse imaging suggests that RGD domain of MMP9 FNII domain anchors integrins at the cell membrane.

Recent research has demonstrated that MDA-MB-231 cells secrete MMP9-enriched exosomes, which can diffuse and degrade the surrounding matrix [57]. Since exosome bio-genesis involves the double invagination of the plasma membrane to form intraluminal bodies (ILB) inside a single multivesicular body (MVB) [58], it is likely that MMP9 is present both in the MVBs which merge with the cell membrane, and is also packaged in the exosomes which get secreted. While the MVB-associated MMPs stabilize ITGβ1 on the cell membrane via the RGD domain, upon matrix degradation mediated by exosomal MMPs, membrane-stabilized ITGβ1 forms firm adhesions with the underlying matrix. However, the details of this mechanism is yet to be established.

In conclusion, our data illustrates the importance of MMP9/ITGβ1 association in sustaining cancer invasion via ITGβ1/MMP9 co-trafficking and ITGβ1 stabilization at the cell membrane and subsequent MMP mediated matrix degradation.

## Acknowledgements

We acknowledge IIT Bombay for providing Bio-AFM, FACS and Confocal Microscopy facilities.

## Author Contributions

Conceptualization: S.D., S.Sen; Methodology: S.D., S.S.; Formal analysis: S.D., S.T., M.A.H; Investigation: S.D., S.Sen.; Data curation: S.D.; Writing - original draft: S.D., S.Sen; Writing - review & editing: S.Sen; Supervision: S.Sen; Project administration: S.Sen; Funding acquisition: S.Sen

