## Supplementary Data for "Two distinct integrin binding sites on MMP9 drive cancer invasion by mediating integrin membrane trafficking & stabilization"

**Supplementary Table 1:**

| **Primer Name** | **Primer Sequences** |
|---|---|
| M9_F_KpnI | CGGGGTACCCCGTCTGCCCTCACCATGAGCCTCTG |
| M9_R_NotI | ATTTGCGGCCGCTTTAAGTCCTCAGGGCACTGCAGGATG |
| M9_F_EcoRI | GCGAATTCCTCACCATGAGCCTCTGGC |
| eGFP_R_XbaI | CGTCTAGACTTGTACAGCTCGTCCATGCCG |
| M9_E402A_F | GCGGCGCATGCGTTCGGCCAC |
| M9_E402A_R | CACGAGGAACAAACTGTATCCTTGGTC |
| M9_RGE_F | GCCGCGGAGAAGGGCGCCTCT |
| M9_RGE_R | CCTCGCTGGTACAGGTCGAGTACTC |
| M9_ΔB1_F | CAGGACCGCTTCTACTGG |
| M9_ΔB1_R | CATCCGGTCCACCTCGCT |
| CycA-F | TGGGCCGCGTCTCCTTTGA |
| CycA-R | GGACTTGCCACCAGTGCCATTA |
| MMP9-F | ACGACGTCTTCCAGTACCGAG |
| MMP9-R | GTAGCCCACTTGGTCCACCT |
| MMP2_HF | AGTGGATGCCGCCTTTAACTG |
| MMP2_HR | GAAGAAGTAGCTGTGACCGCC |
| MMP14_HF | TCAAGTTCAGTGCCTACCGAAGAC |
| MMP14_HR | CAGGTAGCCATATTGCTGTAGCCA |
| MMP1_HF | CTGCTTACGAATTTGCCGACAGAG |
| MMP1_HR | ACAGTTCTAGGGAAGCCAAAGGAG |

**Supp. Figure Legends**

**Supplementary Figure 1**

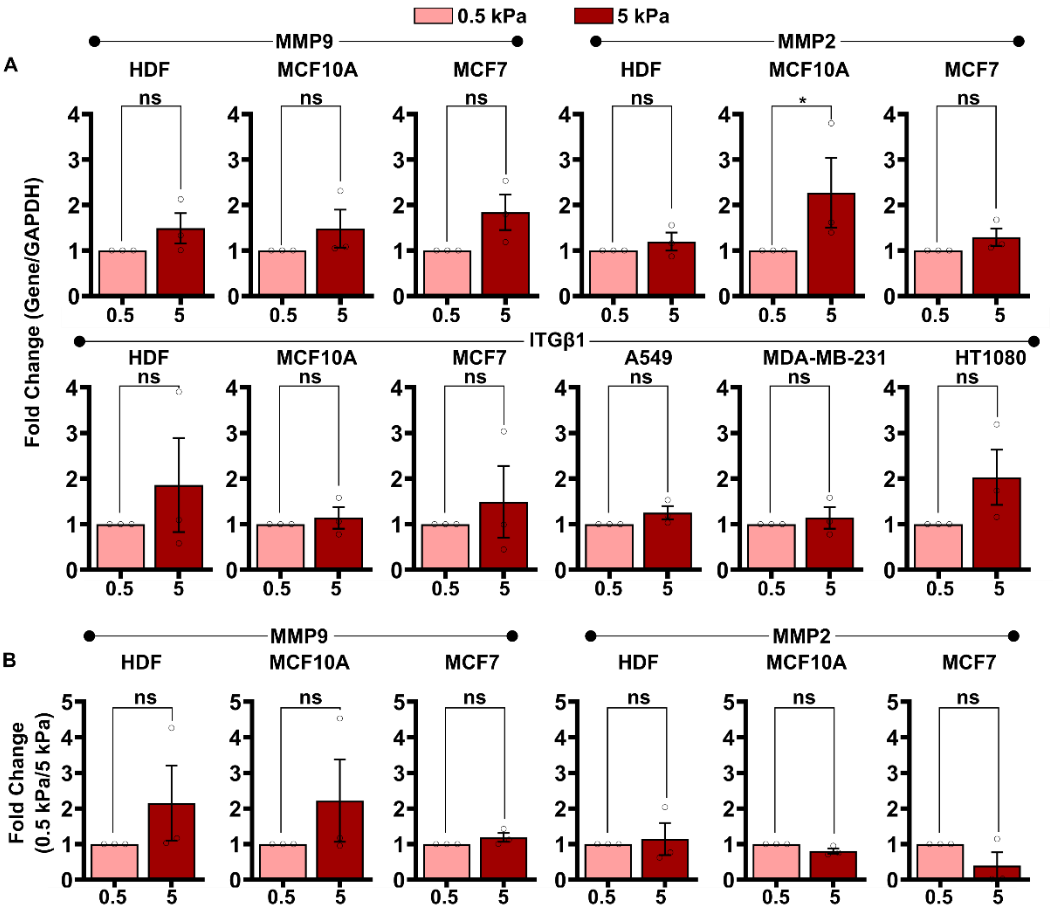


Fold change analysis of MMP2/9 and ITGβ1 expression in HDF, MCF10A, MCF7, A549, MDA-MB-231, and HT1080 cells on 5 kPa gels compared with 0.5 kPa gels from (A) western blot and (B) gelatin zymography. The error bars represent the densitometric analysis from the three independent blots/zymograms ($N = 3$).$* p < 0.05$, significance was measured by the Mann-Whitney U test.

**Supplementary Figure 2:**

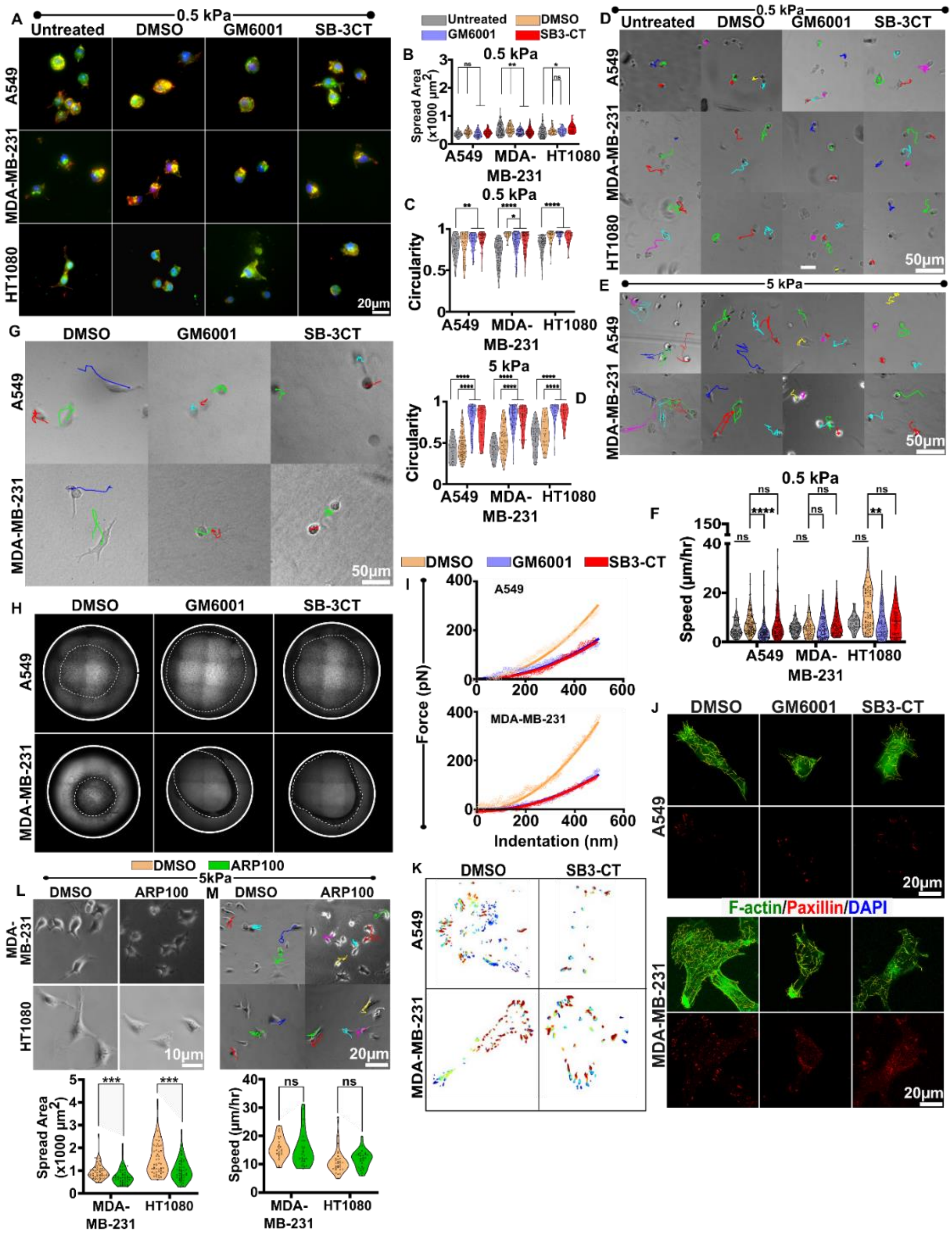


(A) Representative images of Lectin (green), F-actin (red) and DAPI (blue) stained A549, MDA-MB-231, and HT1080 cells on 0.5 kPa gels treated with DMSO, GM6001 and SB3-CT with untreated cells serving as control. (B, C) Quantification of cell spread area and cell circularity ($\approx 60-80$ cells per condition, $N=3$). (D, E) Migration trajectories of cells on 0.5 kPa and 5 kPa gels. (F) Quantification of the 2D motility speed of the cells on 0.5 kPa gels ($\approx 50-60$ cells per condition, $N=3$). (G) Invasion trajectories of DMSO, GM6001 and SB3-CT treated MDA-MB-231 and A549 cells inside 3D collagen gels. (H) Representative images of gel compaction assay of MDA-MB-231 and A549 cells (solid line represents well boundary and dotted line represents final gel boundary). (I) AFM Force curve of MDA-MB-231 and A549 cells. (J) Representative images of F-actin (green) and paxillin (red) stained DMSO, GM6001 and SB3-CT treated MDA-MB-231 and A549 cells. (K) Paxillin adhesion lifetime heatmap of A549 and MDA-MB-231 cells treated with DMSO and SB3-CT. (L) Representative images of DMSO/APR100 treated MDA-MB-231 and HT1080 cells and quantification of cell spread area ($\approx 40-50$ cells per condition, $N=3$). (M) Representative migration trajectories of DMSO/APR100 treated MDA-MB-231 and HT1080 cells on 5 kPa gels and quantification of cell speed ($\approx 40-60$ cells per condition, $N=3$). Significance was measured by One-Way ANOVA or nonparametric test ($* p<0.05, ** p<0.01, *** p<0.001, **** p<0.0001$).

**Supplementary Figure 3:**

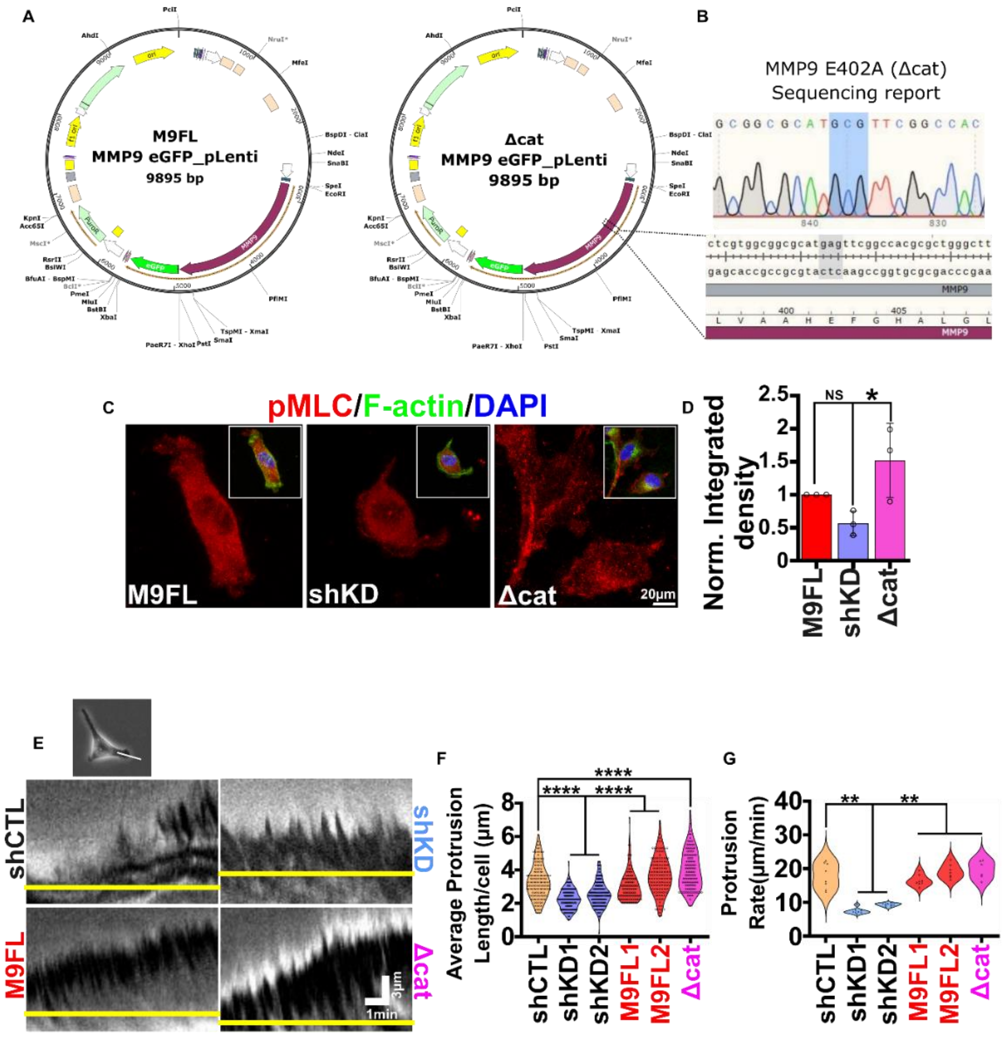


(A) eGFP fused M9FL and Δcat lentiviral pLenti plasmids and (B) sequence verification Δcat cDNA. (C, D) Representative pMLC (red), F-actin (green) and DAPI (blue) stained (inset: merged) images of M9FL, shKD and Δcat HT1080 cells and quantification of pMLC intensity ($\approx 50-60$ cells per condition, $N = 3$). (E-G) Representative kymograms of shCTL, shKD, M9FL and Δcat cells and quantification of the protrusion length ($\approx 50-60$ cells per condition, $N = 3$) and protrusion rate ($\approx 10-15$ cells per condition, $N = 3$). Significance was measured by One-Way ANOVA or nonparametric test ($* p < 0.05, ** p < 0.01, *** p < 0.001, **** p < 0.0001$).

**Supplementary Figure 4:**

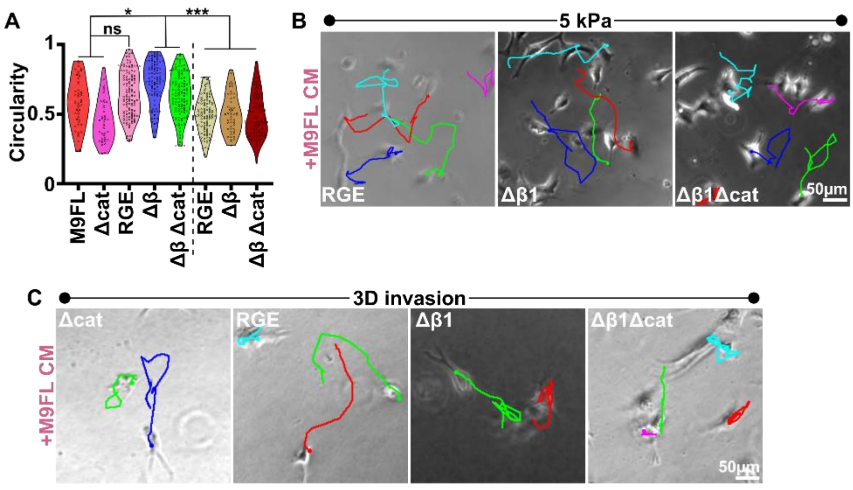


(A) Circularity of M9FL, Δcat, RGE, Δβ1, and Δβ1Δcat cells ($\approx 70-80$ cells per condition, $N = 3$). (B) Representative trajectories of 2D migration on 5 kPa gels of RGE, Δβ1, and Δβ1Δcat cells supplemented with conditioned media from M9FL. (C) 3D invasion trajectories of Δcat, RGE, Δβ1, and Δβ1Δcat inside 3D collagen supplemented with conditioned media from M9FL.

**Supplementary Figure 5:**

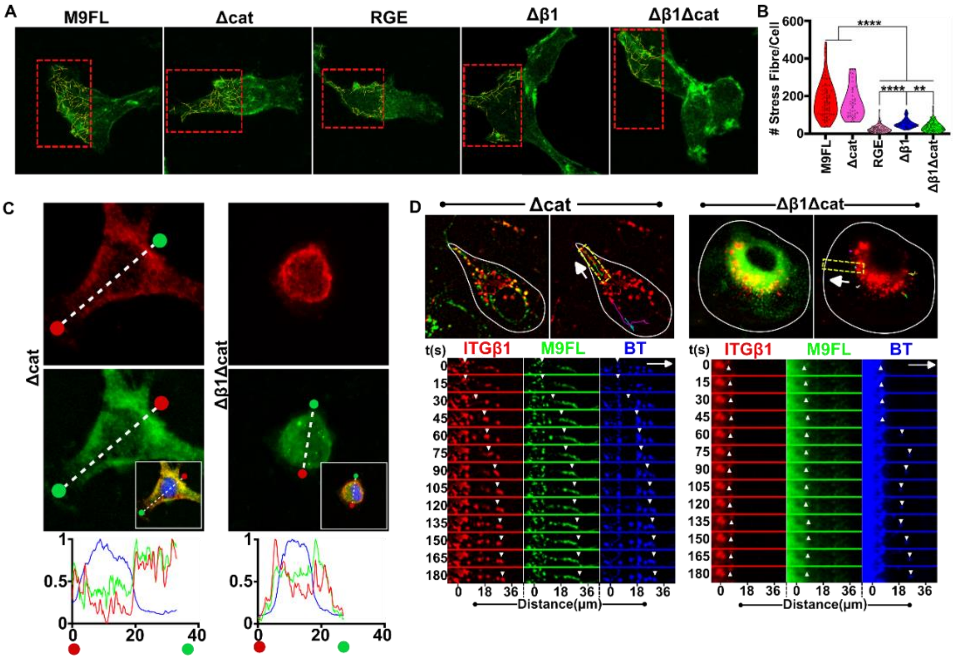


(A) Representation of F-actin (green) stained M9FL, Δcat, RGE, Δβ1, and Δβ1Δcat cells. Red dotted highlighted area represents processed stress fibres (yellow lines). (B) Quantification of the number of stress fibres per cell was done ($\approx 40 - 50$ cells per condition, $N = 3$). (C) Representative images of ITGβ1 (red), MMP9 (green), and DAPI (blue) stained Δcat and Δβ1Δcat cells, with intensity along the Line profiles drawn in the direction of cell migration (magenta and yellow circles depict the migration front and cell rear respectively). (D) Trajectories of ITGβ1 (red) foci in Δcat and Δβ1Δcat cells along the direction of migration (white arrow). Merged images of MMP9 (green) and ITGβ1 foci (red) are also shown. Timestamp images of MMP9 (green), ITGβ1 foci (red) and bio-tracker (blue) of the indicated dotted box (inset) are represented.

**Supp. Movie Legends**

Supplementary Movie 1: Timelapse images of A549, MDA, and HT cells cultured on top of 0.5 kPa PA gels in the presence of DMSO, GM6001 and SB3-CT. Untreated cells served as control.

Supplementary Movie 2: Timelapse images of A549, MDA, and HT cells cultured on top of 5 kPa PA gels in the presence of DMSO, GM6001 and SB3-CT. Untreated cells served as control.

Supplementary Movie 3: Timelapse images of A549, MDA, and HT cells cultured inside 3D collagen gels in the presence of DMSO, GM6001 and SB3-CT.

Supplementary Movie 4: Timelapse images of the focal adhesion turnover of DMSO and SB3-CT treated A549, MDA, and HT cells transfected with paxillin eGFP.

Supplementary Movie 5: Timelapse images of DMSO/ARP100 treated MDA and HT cells cultured on 5 kPa PA gels.

Supplementary Movie 6: Timelapse images of HT1080 shCTL, shKD, M9FL and Δcat cells cultured on 5 kPa PA gels.

Supplementary Movie 7: Timelapse images of HT1080 shCTL, shKD, M9FL,and Δcat cells inside 3D collagen gels.

Supplementary Movie 8: Timelapse images of the focal adhesion turnover of HT1080 M9FL, shKD and Δcat cells transfected with paxillin eGFP plasmid.

Supplementary Movie 9: Timelapse images of HT1080 M9FL, Δcat, RGE, Δβ1 and Δβ1Δcat cells cultured on 5 kPa PA gels.

Supplementary Movie 10: Timelapse images of HT1080 RGE, Δβ1 and Δβ1Δcat cells supplemented with conditioned media from HT1080 M9FL cells on top of 5 kPa gel gels.

Supplementary Movie 11: Timelapse images of HT1080 M9FL, Δcat, RGE, Δβ1 and Δβ1Δcat cells inside 3D collagen gels.

Supplementary Movie 12: Timelapse images of HT1080 RGE, Δβ1 and, Δβ1Δcat cells supplemented with conditioned media from HT1080 M9FL cells inside 3D collagen gels.

Supplementary Movie 13: Timelapse images of focal adhesion turnover of HT1080 M9FL, Δcat, RGE, Δβ1 and Δβ1Δcat cells transfected with paxillin eGFP plasmid.

Supplementary Movie 14-18: Timelapse images of M9FL (Supp. Movie 14), Δcat (Supp. Movie 15), RGE (Supp. Movie 16), Δβ1 (Supp. Movie 17) and, Δβ1Δcat cells (Supp. Movie 18) (MMP9-eGFP expressing) transfected with ITGβ1-mRFP plasmid (red) and stained with bio-tracker (blue). The white line represents the cell boundary while the arrow represents the direction of cell migration.
